# The co-stimulation of anti-CD28 and IL-2 enhances the sensitivity of ELISPOT assays for detection of neoantigen-specific T cells in PBMC

**DOI:** 10.1101/760355

**Authors:** Yunxia Tang, Linnan Zhu, Qumiao Xu, Xiuqing Zhang, Bo Li, Leo J. Lee

## Abstract

Neoantigen-based cancer immunotherapies hold the promise of being a truly personalized, effective treatment for diverse cancer types. ELISPOT assays, as a powerful experimental technique, can verify the existence of antigen specific T cells to support basic clinical research and monitor clinical trials. However, despite the high sensitivity of ELISPOT assays, detecting immune responses of neoantigen specific T cells in a patient or healthy donor’s PBMCs is still extremely difficult, since the frequency of these T cells can be very low. We developed a novel experimental method, by co-stimulation of T cells with anti-CD28 and IL-2 at the beginning of ELISPOT, to further increase the sensitivity of ELISPOT and mitigate the challenge introduced by low frequency T cells. Under the optimal concentration of 1μg/mL for anti-CD28 and 1U/ml for IL-2, our new method can increase sensitivity by up to 5 folds comparing to a conventional ELISPOT, and outperforms other cytokine stimulation alternatives. To the best of our knowledge, this is the first report that the co-stimulation of anti-CD28 and IL-2 is able to significantly improve the sensitivity of ELISPOT assays, indicating that anti-CD28 and IL-2 signaling can act in synergy to lower the T cell activation threshold and trigger more neoantigen-specific T cells.

---

Neoantigens are tumor-specific antigens derived from somatic mutations in cancer cells, and it has been shown that the existence of neoantigen specific T cells could contribute to tumor shrinkage[1], impact the effectiveness of immune checkpoint blockade therapies[2, 3], and lead to novel treatment strategies such as personalized cancer vaccines[4, 5]. Experimentally, the detection of antigen-specific T-cells can be conducted by several different assays, including cytokine enzyme-linked immune spot assay (ELISPOT), staining with tetrameric major histocompatibility complex (MHC)/peptide complexes and intracellular cytokine staining (ICS) etc. Staining with tetramers can only detect binding between TCR and MHC/peptide complexes, but not IFN-γ response of T cells. ICS requires knowledge of both MHC alleles and the dominant T-cell epitopes within the protein of interest, which is time consuming to obtain and needs a large number of T cells (10^6) to start with. Ex vivo ELISPOT is a well-established method for the assessment of the functionality of antigen-specific T cells [6]. It only requires a relatively low number of T cells (0.1-1×10^5) and can detect peptide specific T cell responses with simplicity and speed. ELISPOT assays are generally considered to be a highly sensitive method, which can detect one in 10,000 cells. However, neoantigen specific CD8+ TCR frequencies can be as low as 0.004% of total PBMC [7], and a more sensitive method than standard ELISPOT is needed to detect the IFN-γ cytokine produced by neoantigen specific CD8+ T cells.

Several approaches that can improve the sensitivity of ELISPOT assays have been reported. The most widely accepted one is pre-stimulation of PBMC with antigens. It has been demonstrated [8–10] that antigen-specific T cell responses can be amplified by culturing T cells from PBMCs with peptide for ~10 days prior to performing ELISPOT. T cell responses detected by the above method are 2-5 folds greater than standard ELISPOT assays performed overnight on PBMCs *ex vivo.* However, this method is time and labor consuming. Mallone et al found that addition of low dose IL-2 led to a consistent improvement in ELISPOT sensitivity, as assessed by measuring low grade CD8+ T cell responses against β-cell autoantigens in T1D patients[11]. Martinuzzi et al[12] found that addition of low dose IL-7 in ELISPOT proved capable of amplifying the low-grade CD8+ T cell responses against β-cell epitopes (1.5-fold increase). Ott et al [13] conducted ELISPOT assays to measure T cell responses with and without the addition of anti-CD28 antibody. A 3.9-fold increase of T cell response was observed, and sometimes IFN-γ responses could only be detected when anti-CD28 antibody was present. Calarota et al[14] evaluated weather the IFN-γ production by antigen specific stimulated lymphocytes could be increased by IL-15 in rhesus macaques infected with SIVmac251, and showed that it significantly increased IFN-γ production, with mean IFN-γ spot number increasing of 2.5 and 1.8 folds in response to SIV gag and env peptide pools, respectively. Jennes et al [15] reported that addition of IL-7 and IL-15 increased the number of PPD-specific CD4+ T cells up to 2.4-fold in fresh PBMC. Since these modified ELISPOT assays were useful in enhancing the sensitivity of T cell responses to viral or autoimmune antigens, they could be applied to improve the detection of IFN-γ produced by neoantigen specific T cells as well.

Mechanistically, these cytokines or co-stimulatory molecules play their stimulatory roles through different pathways. IFN-γ secreted by antigen specific T cells depends on two signals. Signal 1 consists of the TCR/MHC-peptide interaction, CD4 and CD8 coreceptors and adhesion molecules. Signal 2 is mediated by co-stimulation of a separate set of molecules including CD28. Together, signal 1 and 2 initiate a signal transduction cascade that activates subsequent transcription factors and cytokines to regulate T cell proliferation and differentiation[16]. IL-2 is important for the expansion and survival of Ag-reactive T cells, both for CD4 and CD8 T cells[17]. The magnitude and duration of IL-2 signals can have a profound influence on effector and memory T cells [18, 19], and it can also effectively support the cytotoxic T lymphocyte generation from prevailingly naive CD8+ T cells [20] [21]. Most T cells express CD28, so anti-CD28 can activate T cells including naive T cells. Taken together, the co-stimulation[22] of CD28 and IL-2 signals may boost naïve responses as well as memory/effector ones. In contrast, IL-7 and IL-15 mostly exert their effects on effector or memory T cells [23] [24] [25] [26] [27]. Since naïve T cells is an important source of neoantigen-specific T cells, we set out to test the effect of anti-CD28 plus IL-2 co-stimulation in ELISPOT in this work.

## Material and Methods

### 1 healthy donors

Healthy donors were volunteers recruited from our lab. All healthy donors’ HLA typing include HLA-A*0201.

### 2 peptides

The peptides we used are the following: cytomegalovirus (CMV) pp65 495–503 (NLVPMVATV), and Melan A (EAAGIGILTV). PHA was used as positive control (5 μg/mL; Sigma).

### 3 neoantigen

To identify neoantigens, we used a neoantigen prediction algorism to identify which mutated peptides in cancer cells could be present on the cell surface by HLA molecules. Using this inhouse pipeline [28], we analyzed 60 neoantigen peptides presented by HLA-A*0201 molecules across different cancer types from the TCGA cohort. The affinity of the neoantigens were verified with an affinity test by T2 cells. Thirty neoantigen peptides of high affinity with HLA-A*0201 were selected. Neoantigens were synthesized and HPLC purified by Genscript Inc (Nanjing, Jiangsu, China) to greater than 98% purity. The sequence of the neoantigen peptides used in this work were

#50 ILCATYVKV
#51 GLKDLLNPI
#61 ILVDWLFEV
#72 YLILWCFYL
#74 KLMGIVYKV
#91 FLDPALYPL
#1 VLNCLLYAV
#2 GIMESFFTV

### 4 Pre-stimulation of PBMC

CD8+ T cells from PBMC were enriched with CD8+ T-cell isolation kit beads (Miltenyi Biotech), then they were frozen until DC is mature. For in vitro pre-stimulation of antigen-specific T cells, CD8+T cells were thawed and cultured in AIM-V medium supplemented with 10% autologous serum. The cells were stimulated in 24-well cell culture plates at 5×10^6 cells per well with autologous DC pulsed with individual antigen (50μg/ mL) in the presence of IL-7 (20 ng/mL; Pepro tech). On day 3, IL-2 (20 U/mL; Pepro tech) was added. Half-medium change and supplementation of cytokines were performed every 3 days, as described previously [4]. After 10 days, T-cell specificity was tested against peptide by IFN-γ ELISPOT assays.

DC were cultured by following the fast DC procedure[29]. PBMC were isolated from peripheral blood of healthy donors by Ficoll gradient centrifugation. The MACS CD14 isolation kit (Miltenyi Biotech, Bergisch Gladbeck, Germany) was used to purify monocytes. CD14 monocytes were subsequently cultured in 24-well plates (10^6 cells/well) in fresh complete medium supplemented with 1000 U/mL GM-CSF and 500 U/mL IL-4 for 24 hours, followed by incubation with the proinflammatory mediators (1000 U/mL TNF-α, 10 ng/mL IL-1β, 10 ng/mL IL-6, and 1 μM PGE_2_) for another 24 hours.

### 5 ELISPOT assay

Human IFN-γ ELISPOT assays were performed by ELISPOT kit (Mab Tech). The plates were washed three times with PBS. PBMC were then added to individual wells in AIM-V in the presence or absence of anti-CD28 antibody (Biolegend), IL-2, IL-7 and IL-15 (Pepro tech) at concentrations specified in the Results section. The plated cell number was 3 ×10^5 PBMC per well. Antigens were added at previously established optimal concentrations for PBMC stimulation in the ELISPOT assays[30]: CMV protein antigen 50μg/mL, Melan A antigen 50μg/mL, Phytohemagglutinin (PHA, Sigma) 5μg/mL. The cells were incubated at 37°C for 18-24 h in the presence of 5% CO_2_ and then washed five times with PBS. The detection antibody, 7-B6-ALP was diluted 1:200 in PBS (0.5% FBS), 100μL per well. After incubation at 37°C for 2 h, plates were washed five times with PBS and incubated for 10 min at room temperature with BCIP/NBT- plus substrate. The resulting spots were counted using a computer-assisted ELISPOT image analyzer (Biosys bioreader 4000) and custom software were designed to detect spots using predetermined criteria based on size, shape, and colorimetric density [15]. The measurement of spot-size distribution is a built-in function of the software. Antigen-specific frequencies of cytokine producing cells were calculated by the number of detected spots obtained in the presence of relevant antigen (performed in duplicate). Following the established guideline [30], a positive response was defined when the mean of the antigen-stimulated replicates was greater than or equal to ten spots per well, and the mean of the antigen- stimulated replicates was greater than three times the mean of the replicates from the negative control wells.

### 6 Statistical analysis

Since spot numbers are normally distributed within each triplicate, comparisons of means within the same donor were carried out with two-tailed Student’s t-test, while the non-parametric Wilcoxon matched pairs test was used for comparing two ELISPOT conditions across the whole cohort of donors. Analysis of variance (ANOVA) was used for comparisons among multiple groups.

## Results

### Optimal concentrations of anti-CD28 and IL-2 are 1μg/mL and 1U/mL

According to previous reports, 0.5 U/ml IL-2 [11] or 1μg/mL anti-CD28 [13] can improve the sensitivity of ELISPOT. Can the sensitivity be further improved when these two factors are combined? To answer such a question, we performed some preliminary experiments by adding both IL-2 (0.5 U/ml) and anti-CD28 (1μg/mL) for co-stimulation, and the results are shown in Fig.1A. When we stimulated PBMC with CMV peptide, the number of IFN-γ spot increased by 2 times compared with anti-CD28 alone at 1μg/mL, or 2.6 times compared with IL-2 alone at 0.5 U/ml. When we stimulated PBMC with Melan A peptide, the number of IFN-γ spot increased by 2.2 times (compared with anti-CD28 alone) or 1.8 times (compared with IL-2 alone). Meanwhile, the spot numbers of the control group (PBMC+DMSO) stayed roughly the same. It is clear that the combination of these two factors can further improve the sensitivity of ELISPOT.

Next, in order to find out the optimal concentrations of anti-CD28 and IL-2 co-stimulation for PBMC ELISPOT assays, different concentrations of these two stimulators were tested systematically. We first fixed the concentration of IL-2 to 0.5 u/ml and adjusted the concentration of anti-CD28. As shown in Fig. 1B, for PBMC cocultured with both CMV and Melan A peptides, with the concentrations of anti-CD28 increasing from 0.25μg/mL to 2μg /mL, the mean IFN-γ spot numbers also increased constantly. Ideally, we would like to have the mean IFN-γ spot numbers of the control group (PBMC plus DMSO) stay constant with varying anti-CD28 concentrations. What was actually observed was that it stayed almost constant for anti-CD28 concentrations of 0, 0.25, 0.5 and 1μg /mL (13, 16, 20 and 23, respectively) but a sharp increase from 23 to 70 when the concentration of anti-CD28 reaches 2μg/mL. Therefore, the best concentration of anti-CD28 is 1μg /mL. Next, we fixed the concentration of anti-CD28 to 1μg /mL and varied the concentration of IL-2. As shown in Fig. 1C, for both CMV and Melan A peptides, the average IFN-γ spot numbers increased constantly with 0, 0.5, 1, 5 and 10U/mL of IL-2 stimulation, while the IFN-γ spot numbers of PBMC plus DMSO increased significantly from 19 to 55 when the concentration of IL-2 was increased from 1U/mL to 5U/mL. We also performed an extra experiment with another volunteer’s PMBC, and the IFN-γ spot numbers of the control group were 5, 25, and 30 respectively with 1, 2.5 and 5U/mL of IL-2 stimulation (data not shown). Therefore, the optimal concentration of IL-2 is 1 U/mL. According to the results above, we chose 1 U/mL IL-2 and 1μg /mL anti-CD28 as the optimal concentrations for co-stimulation and used them in the remaining experiments of this work.

**Figure 1.**
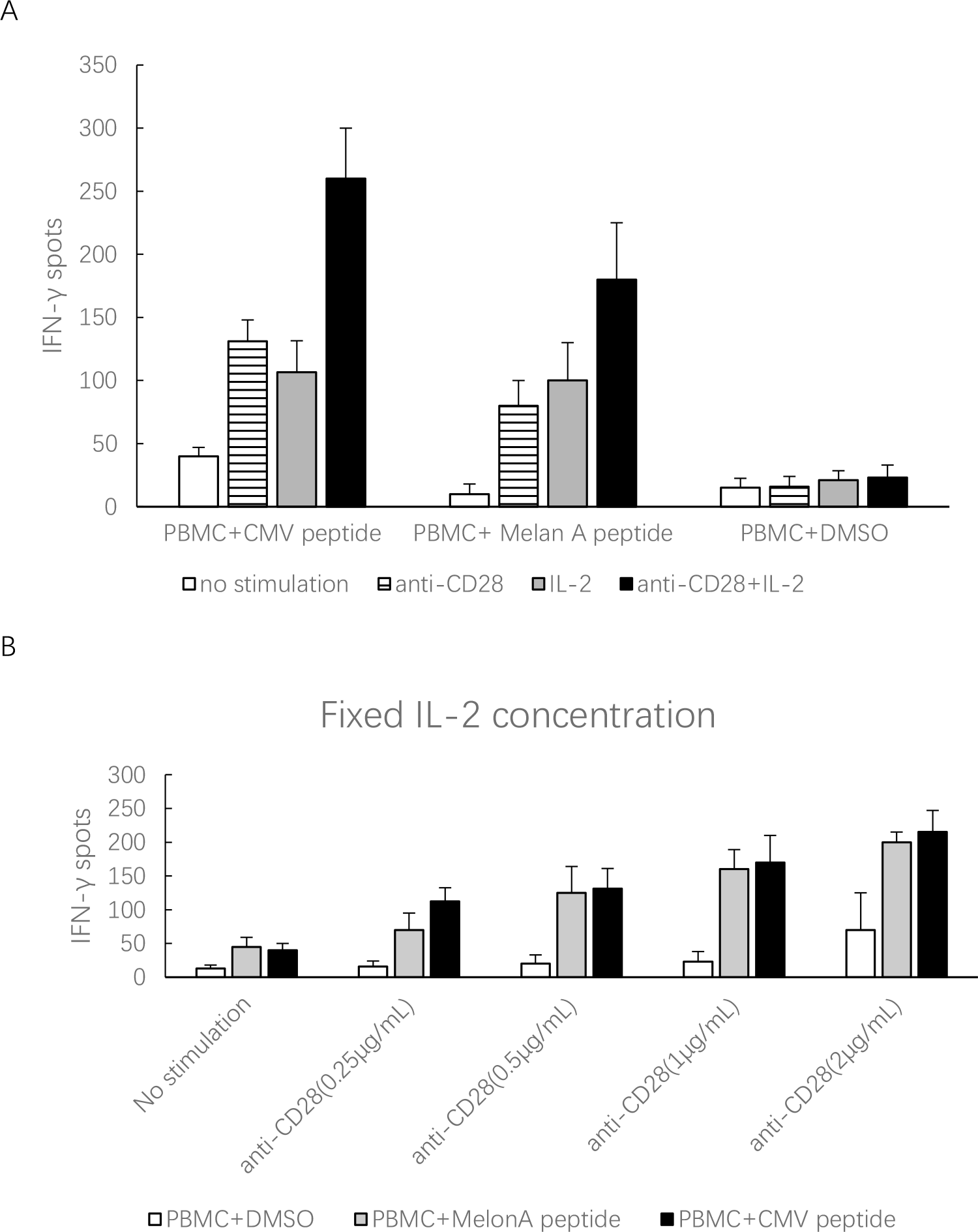
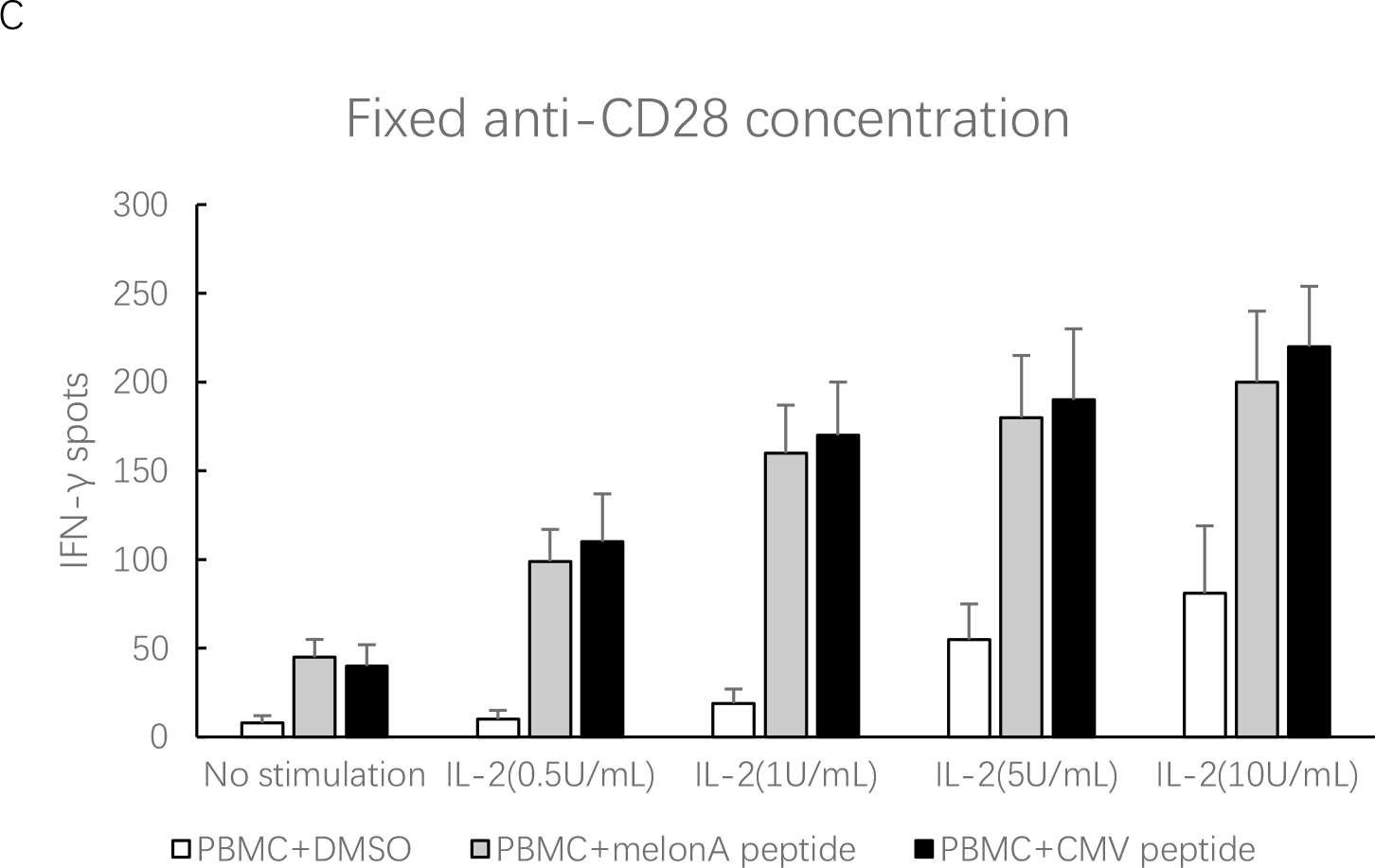
1μg/mL anti-CD28 and 1 U/mL IL-2 are optimal concentrations for co-stimulation. A. IL-2 plus anti-CD28 can improve the sensitivity of ELISPOT compared with IL-2 or anti-CD28 alone. When we stimulated PBMC with CMV peptide, the number of IFN-γ spots increased by 2 times (compared with anti-CD28) or 2.6 times (compared with IL-2). When we stimulated PBMC with Melan A peptide, the number of IFN-γ spots increased by 2.2 times (compared with anti-CD28) or 1.8 times (compared with IL-2).
B. IL-2 concentration was fixed to 0.5 u/ml and anti-CD28 concentration was adjusted from 0.25μg/mL to 2μg /mL. The optimal concentration of anti-CD28 is 1μg/mL when combined with IL-2.
C. Anti-CD28 concentration was fixed to 1μg/mL and IL-2 concentration was adjusted from 0.5U/mL to 10U/mL. The optimal concentration of IL-2 is 1 U/mL when combined with anti-CD28.

### Addition of anti-CD28 plus IL-2 stimulates more IFN-γ producing T cells than IL-7 or IL-15

As previously reported, addition of IL-7 and IL-15 can also enhance the sensitivity of ELISPOT assays in detecting antigen-specific T cells, albeit mostly for memory or effector T cells. To test if the co-stimulation by anti-CD28 and IL-2 can be more effective than IL-7 or IL-15, we further carried out experiments to compare them in the context of PBMC co-cultured with CMV or Melan A peptides, both of which are usually responded by memory or effector T cells. The experiments were performed using three healthy donors’ PBMCs, each with three repeats under every condition. As shown in Fig. 2A, for CMV peptides, the mean IFN-γ spot number when co-stimulated by anti-CD28 plus IL-2 was significantly higher than those of IL-7 or IL-15 alone, in addition to those of anti-CD28 or IL-2 individually. The fold changes across these three volunteers’ PBMCs were summarized in Fig. 2B, where the co-stimulation of anti-CD28 plus IL-2 reached an 11-fold change, while other stimulators were 2-8 folds relative to the control group. Similar results were obtained with Melan A peptides, both in terms of IFN-γ spot numbers (Fig. 2C) and fold changes (Fig. 2D). Typical well pictures of each group were also shown in Fig. 2E. Since IL-7 and IL-15 has little effect on naïve T cells, which is important for neoantigen-specific T cell detection, we didn’t spend the extra effort to test the effectiveness of co-stimulation by both IL-7 and IL-15. In addition, co-stimulation by anti-CD28 and IL-2 is also more cost-effective than IL-7 and IL-15, since both reagents are cheaper.

**Figure 2.**
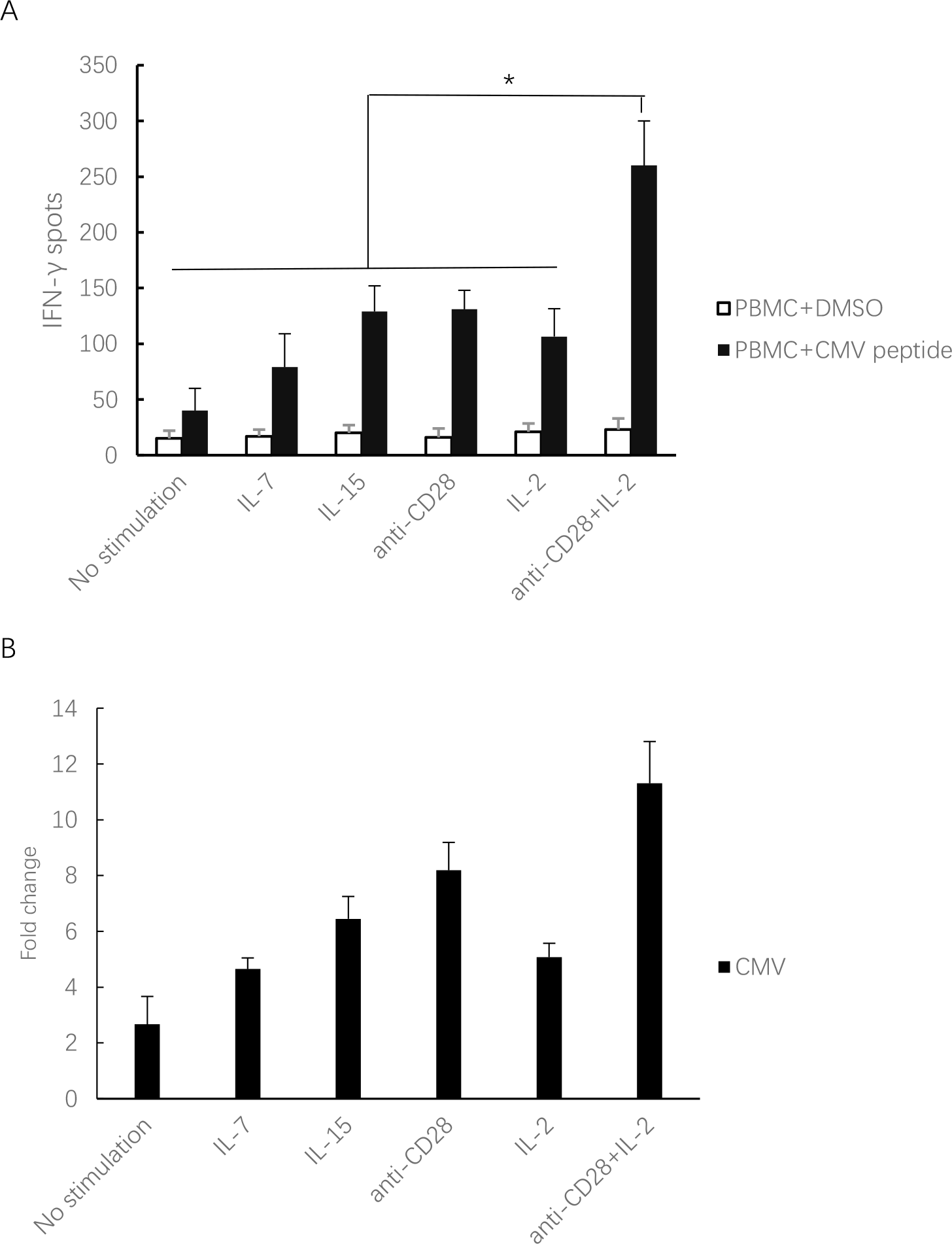
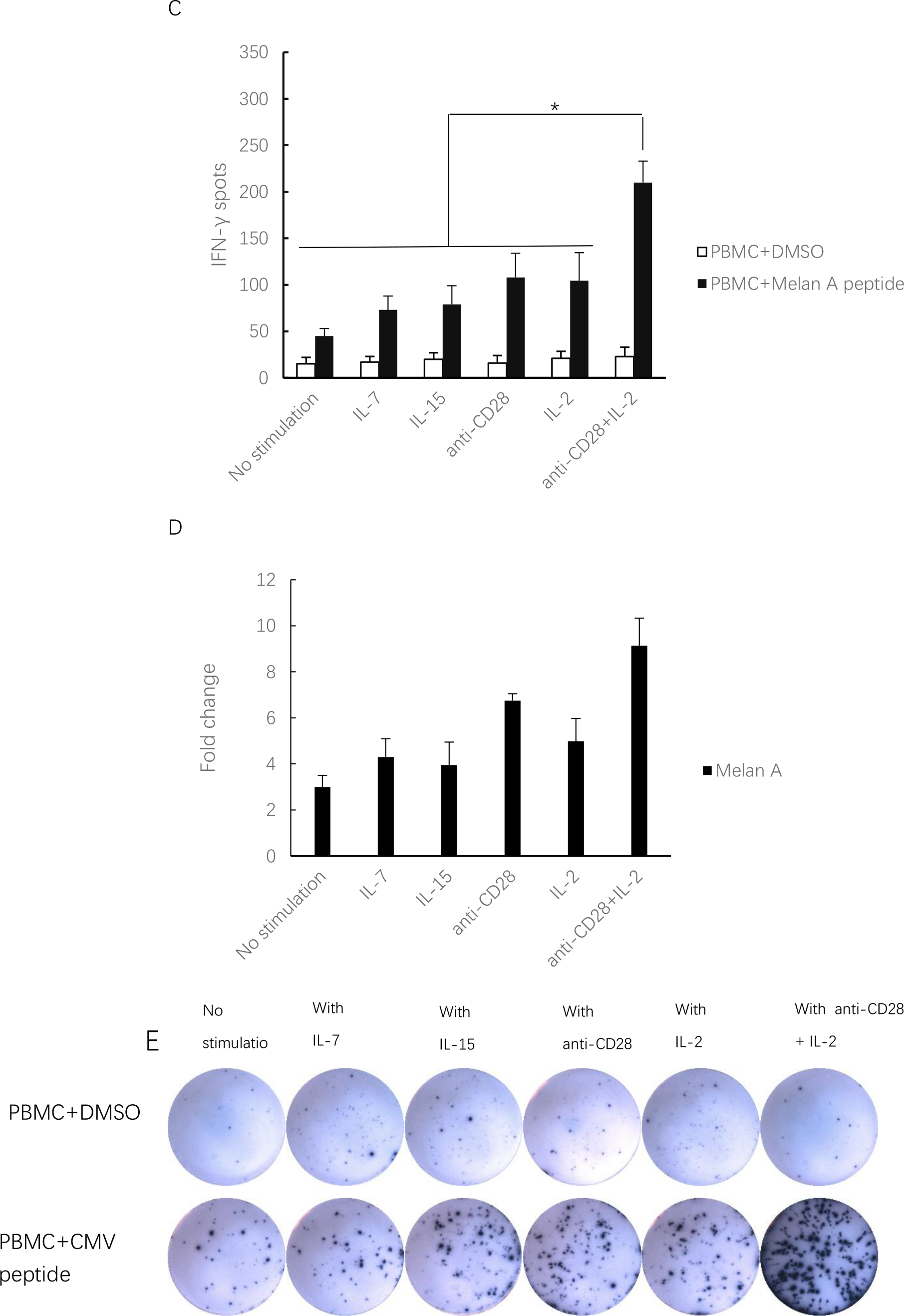
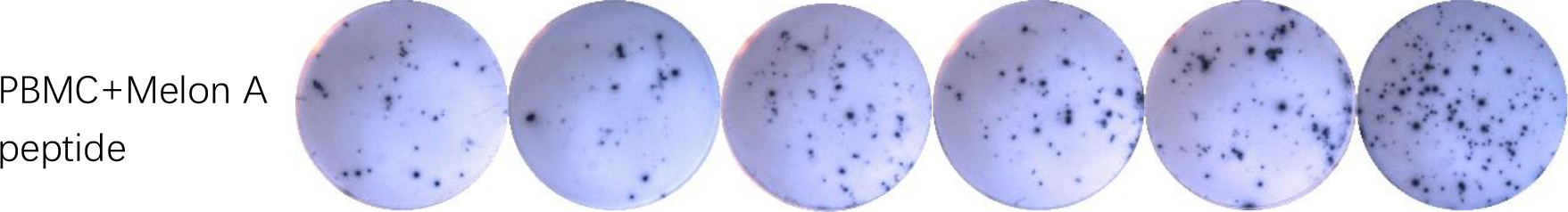
Anti-CD28 and IL-2 co-stimulation enhances the IFN-γ spots of the ELISPOT assays when co-cultured with CMV or Melan A peptide. The experiments were repeated in three independent healthy donors. A. Compared to the group without cytokines, for CMV peptide, more IFN-γ spots were observed in the groups in the presence of IL-7, IL-15, IL-2, anti-CD28 or anti-CD28 plus IL-2. The IFN-γ spot number under the stimulation of anti-CD28 and IL-2 was the highest among all groups. The average IFN-γ spots/300,000 PBMC in the presence of anti-CD28 plus IL-2 was 260, the average IFN-γ spots/300,000 PBMC in the presence of IL-7, IL-15, anti-CD28, IL-2, were 79, 129, 131 and 107.
B. In the case of CMV peptide, IFN-γ spot numbers from IL-7, IL-15, IL-2, anti -CD28 culture were 2-8 times of the control group (PBMC plus DMSO). While the number of anti-CD28 plus IL-2 culture was 11 times of the control group.
C. In the case of Melan A peptide, IFN-γ spot number of anti-CD28 and IL-2 co-stimulation was the highest among all groups. The average IFN-γ spots/300,000 PBMC in the presence of anti-CD28 plus IL-2 was 210, the mean IFN-γ spots in the presence of IL-7, IL-15, anti-CD28, IL-2, were 73, 79, 108 and 105.
D. The average fold change of anti-CD28 plus IL-2 when cultured with Melan A peptide was 9.1, higher than that of other cytokines (3-6 fold change).
E. The spot pictures of ELISPOT assays, one well per group was shown. * indicate P<0.05

### Co-stimulation of anti-CD28 and IL-2 could be applied to neoantigen specific T cell detection in healthy donors’ PBMC

We next tested the performance of co-stimulation by anti-CD28 and IL-2 to detect neoantigen specific T cells in healthy donors’ PBMC, which typically have a very low frequency and contain various types of T cells, especially naive T cells, thus challenging to detect. As detailed in Materials and Methods, SNVs were identified from the TCGA database, while 9-mer peptides containing SNVs with high affinities to HLA-A*0201 were predicted according to PSSMHCpan [28]. We further selected 30 peptides that had been experimentally validated to have high affinities with HLA-A*0201 and tested their immunogenicity using ELISPOT. To establish the ground truth, we first performed ELISPOT assays by 10-day pre-stimulation of T cells from PMBC and neoantigens, a time-consuming but well-accepted method in the field, and a positive response was accepted when the mean of the antigen-stimulated replicates was greater than three folds over the mean of the replicates from the negative control wells. Using a healthy donor’s PBMC, we found 5 positive response neoantigens out of the 30 predicted ones of high affinity, and the IFN-γ spot numbers of these five positive ones (#50, #51, #61, #72, #74) were shown along with three negative ones (#91, #1, #2) in Fig. 3A. We then performed the ELISPOT experiments with co-stimulation of anti-CD28 and IL-2, and detected the same five positive neoantigens (#50, #51, #61, #72, #74) as shown in Fig. 3B. In contrast, under single stimulation conditions (IL-7, IL-15, anti-CD28 or IL-2), only one positive neoantigen (#74) can be detected as having at least 3 times of spot numbers relative to negative control by IL-7 or anti-CD28 (Fig. 3C).

**Figure 3.**
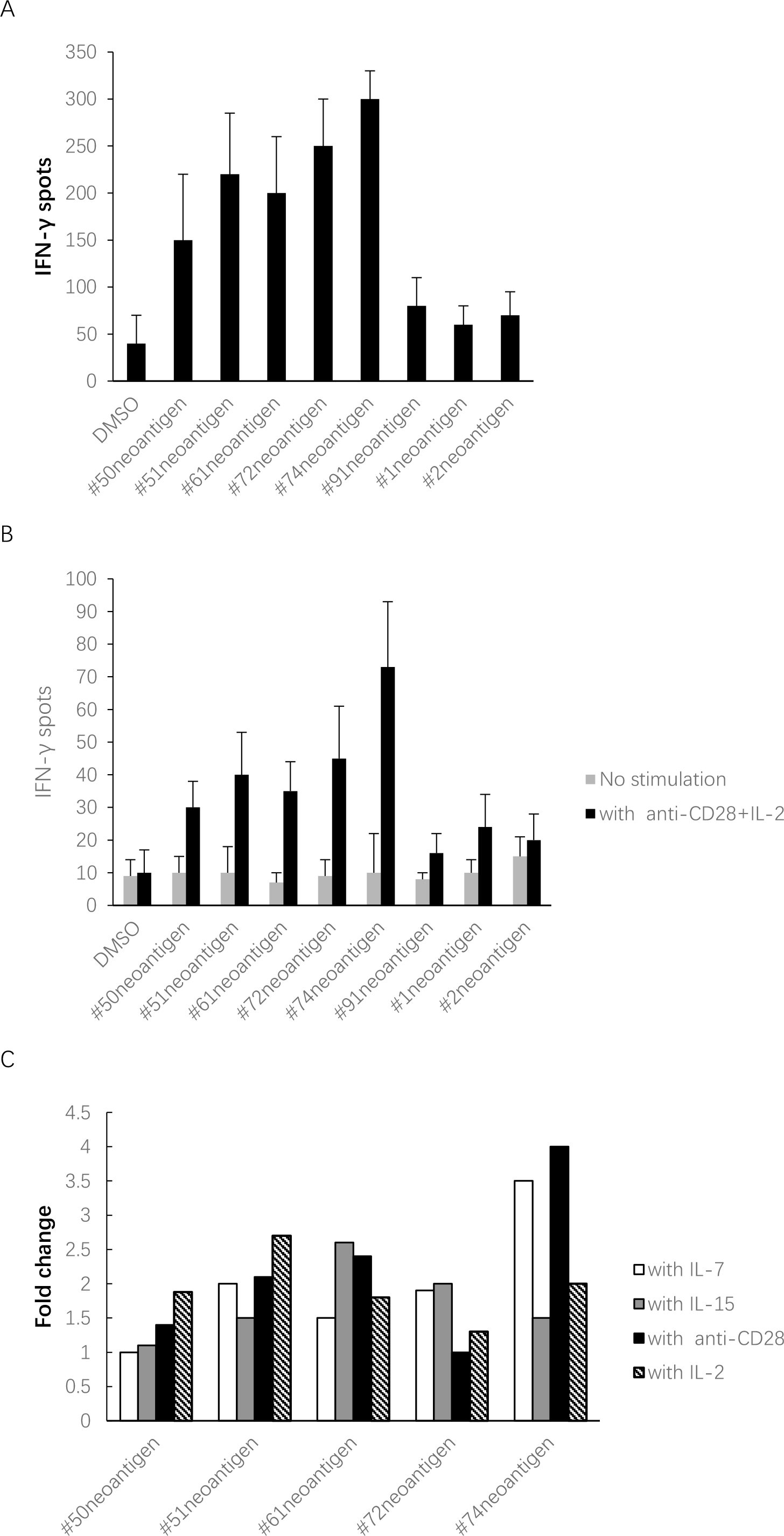
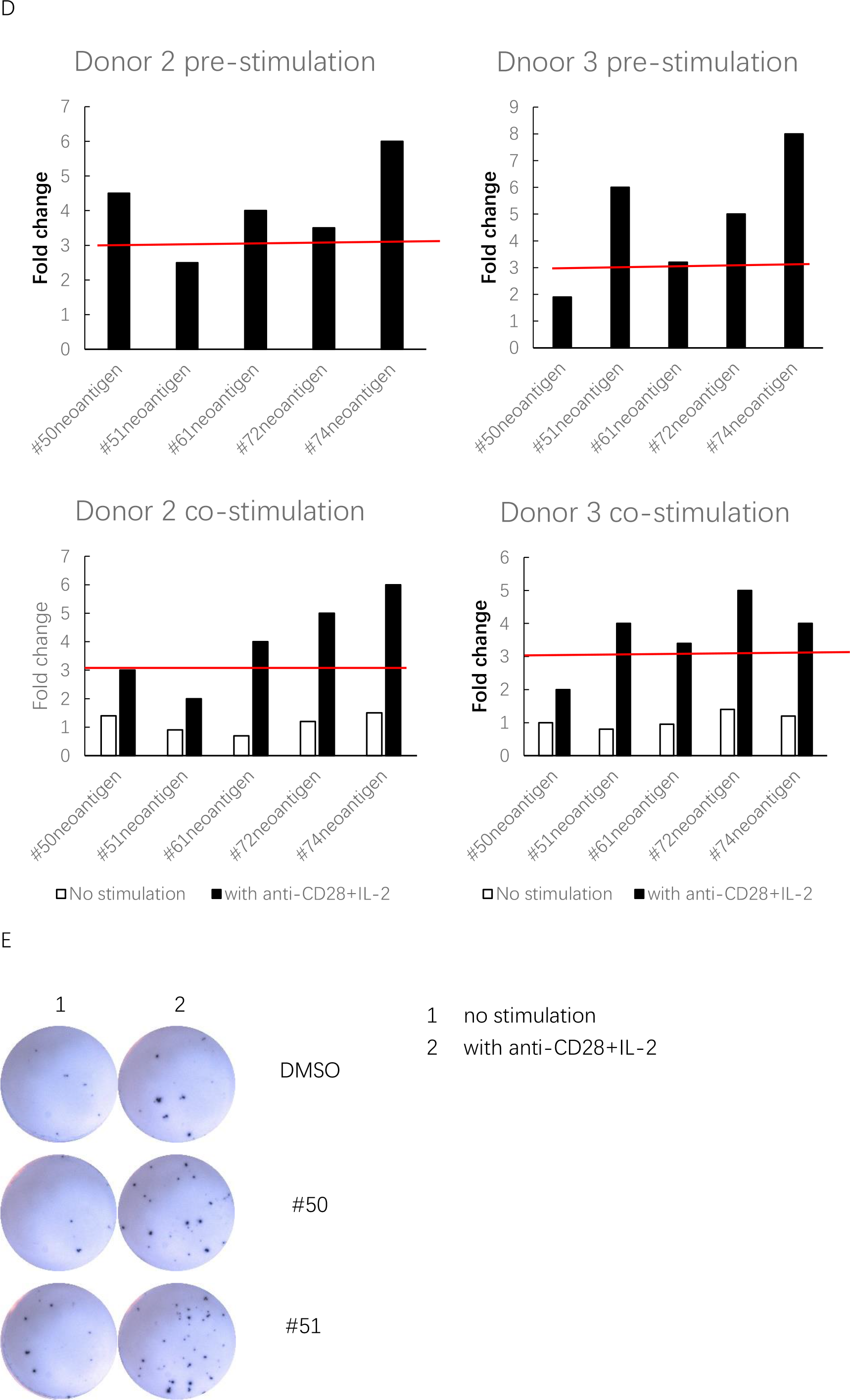
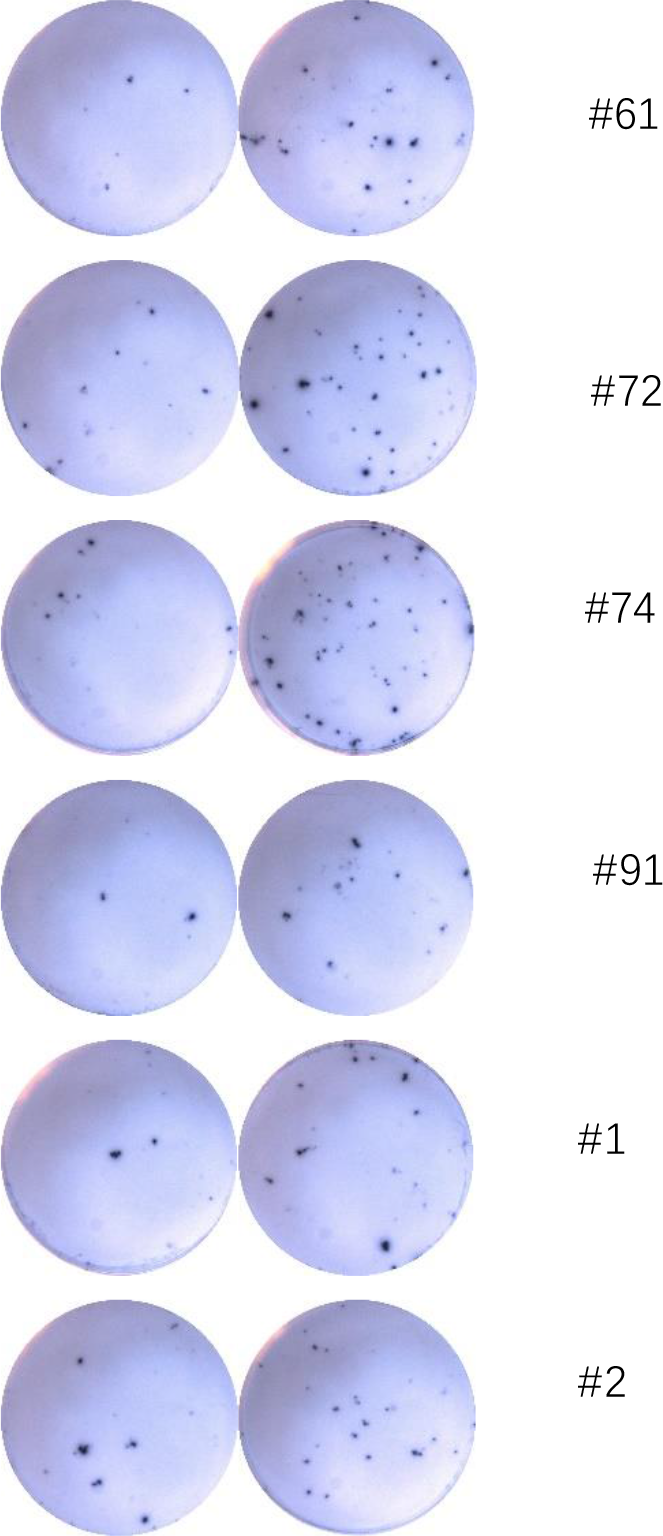
Neoantigen specific T cells can be detected in healthy donors’ PBMC under the co-stimulation of anti-CD28 plus IL-2. A. CD8+T cell responses against #50, #51, #61, #72, #74 neoantigens were detected by pre-stimulation in donor 1, while #91, #1, #2 neoantigen-specific T cells were not detected. The data of other neoantigens which had negative responses were not shown.
B. CD8+T cell responses against #50, #51, #61, #72, #74 neoantigen peptide were detected in the PBMC of donor 1 by co-stimulation of anti-CD28 plus IL-2.
C. The fold changes of IFN-γ spot numbers of different neoantigens with different single cytokine stimulations: only one neoantigen (#74) can be detected, under the stimulation of IL-7 or anti-CD28, as having at least 3 times as many spots as the negative control.
D. Neoantigen-specific T cell detections by pre-stimulation and co-stimulation of anti-CD28 plus IL-2 in donors 2&3: in donor 2, #50, #61, #72, #74 neoantigens had positive responses by pre-stimulation, and the same four neoantigens also had positive responses by co-stimulation; in donor 3, #51, #61, #72, #74 neoantigens had positive responses by pre-stimulation, and the same four neoantigens also had positive responses by co-stimulation.
E. Typical well pictures of ELISPOT assays with donor 1’s PMBC, with or without the co-stimulation of anti-CD28 and IL-2.

We also repeated the same experiments with two more healthy donors’ PMBC (Fig. 3D). In donor 2, we were able to detect four neoantigens (#50, #61, #72, #74) by both pre-stimulation and co-stimulation. In donor 3, we also detected four neoantigens (#51, #61, #72, #74) by both pre-stimulation and co-stimulation. These two different sets of four neoantigens are both subsets of the five detected in donor 1, and the detections between pre-stimulation and co-stimulation were consistent in both cases. However, when only a single stimulator (IL-7, IL-15, anti-CD28 or IL-2) was used, no positive responses were detected in either healthy donors (data not shown). Finally, typical well pictures of donor 1 for eight predicted neoantigens are shown in Fig. 3E. Overall, these results demonstrated both the sensitivity and specificity of our novel co-stimulation approach for detecting neoantigens, comparing with the well-established gold standard of 10-day pre-stimulation.

## Discussion

In summary, we have developed a simple, fast and economic way to significantly enhance the sensitivity of standard ELISPOT assays, and demonstrated its efficacy using different antigens, especially neoantigens discovered from TCGA cancer patients. All our experiments were carried out using PBMCs from healthy donors, which has been previously shown to be a good source of neoantigen-specific T cells, in addition to PBMCs or TILs (tumor infiltrated lymphocytes) from cancer patients. For example, using PBMCs from healthy donors and pre-stimulation, Strønen et al [31] detected neoantigen-specific T cells responding to 11 out of 57 predicted HLA-A2-binding epitopes from three patients. Such an approach was further validated by our results. We also expect our method to work well with cancer patients’ PBMCs, and has the potential to substantially boost previous efforts. For example, by using laborious methods to enrich for neoantigen-specific T cells first, Gros et al [32] identified T cells in three cancer patients’ PBMCs responding to 7 neoantigens with standard ELISPOT, out of a total of 672 predicted ones. In comparison, our simple and fast technique reached a detection rate of 5/60 using PBMCs from a single, WIHLA-type-matched healthy donor, and it could be viewed as a significant improvement.

A major challenge facing neoantigen based personalized cancer immunotherapies is to discover true neoantigens with immunogenicity, which hold the key for effective vaccine or cell-based therapies. Unfortunately, the immunogenicity of a neoantigen is difficult to computationally predict or experimentally validate up to this point. The most reliable way seems to be using a patient’s TIL, which is usually difficult to obtain and culture, and it doesn’t apply to all patients or cancer types. In contrast, PBMCs can be easily obtained from almost all patients, and our novel technique reduced the detection of neoantigen-specific T cells to two days with a small amount of blood, comparing to previous laborious efforts that required weeks or months and a large number of cells. We believe our simple yet effective method is a notable advance, and could pave the way for routine immunogenicity detection/validation for neoantigens in a clinical setting.

## Acknowledgments

This study was supported by National Natural Science Foundation of China (No. S1702926); Science, Technology and Innovation Commission of Shenzhen Municipality under grant No. JCYJ20170303151334808; Shenzhen Municipal Government of China (No. 20170731162715261).

